# Frontotemporal dementia patient-derived iPSC neurons show cell pathological hallmarks and evidence for synaptic dysfunction and DNA damage

**DOI:** 10.1101/2024.04.12.589061

**Authors:** Nadine Huber, Tomi Hietanen, Sami Heikkinen, Anastasia Shakirzyanova, Dorit Hoffmann, Hannah Rostalski, Ashutosh Dhingra, Salvador Rodriguez-Nieto, Sari Kärkkäinen, Marja Koskuvi, Eila Korhonen, Päivi Hartikainen, Katri Pylkäs, Johanna Krüger, Tarja Malm, Mari Takalo, Mikko Hiltunen, Jari Koistinaho, Anne M. Portaankorva, Eino Solje, Annakaisa Haapasalo

## Abstract

Frontotemporal dementia (FTD) is the second most common cause of dementia in patients under 65 years, characterized by diverse clinical symptoms, neuropathologies, and genetic background. Synaptic dysfunction is suggested to play a major role in FTD pathogenesis. Disturbances in the synaptic function can also be associated with the *C9orf72* repeat expansion (C9-HRE), the most common genetic mutation causing FTD. C9-HRE leads to distinct pathological hallmarks, such as *C9orf72* haploinsufficiency and development of toxic RNA foci and dipeptide repeat proteins (DPRs). FTD patient brains, including those carrying the C9-HRE, are also characterized by neuropathologies involving accumulation of TDP-43 and p62/SQSTM1 proteins. This study utilized induced pluripotent stem cell (iPSC)-derived cortical neurons from C9-HRE-carrying or sporadic FTD patients and healthy control individuals. We report that the iPSC neurons derived from C9-HRE carriers developed typical C9-HRE-associated hallmarks, including RNA foci and DPR accumulation. All FTD neurons demonstrated increased TDP-43 nucleus-to-cytosolic shuttling and p62/SQSTM1 accumulation, and changes in nuclear size and morphology. In addition, the FTD neurons displayed reduced number and altered morphologies of dendritic spines and significantly altered synaptic function indicated by a decreased response to stimulation with GABA. These structural and functional synaptic disturbances were accompanied by upregulated gene expression in the FTD neurons related to synaptic function, including synaptic signaling, glutamatergic transmission, and pre- and postsynaptic membrane, as compared to control neurons. Pathways involved in DNA repair were significantly downregulated in FTD neurons. Only one gene, *NUPR2,* potentially involved in DNA damage response, was differentially expressed between the sporadic and C9-HRE-carrying FTD neurons. Our results show that the iPSC neurons from FTD patients recapitulate pathological changes of the FTD brain and strongly support the hypothesis of synaptic dysfunction as a crucial contributor to disease pathogenesis in FTD.

## Introduction

Frontotemporal dementia (FTD) is an umbrella term for a group of neurodegenerative disorders characterized by frontotemporal lobar degeneration, distinct neuropsychological characteristics, altered personality and behavior, (behavioral variant frontotemporal dementia, bvFTD), impairment of speech and language comprehension (primary progressive aphasias, PPAs), and/or disturbances in motor function [1, 2]. FTD is the second most common type of dementia in individuals below 65 years of age [3] and it shares clinical, pathological, molecular, and genetic features with amyotrophic lateral sclerosis (ALS) [4], forming an FTD-ALS disease spectrum. The most common genetic cause of these diseases is the hexanucleotide repeat expansion in the *C9orf72* gene (C9-HRE) [5–9]. Three potentially overlapping toxic mechanisms underlying the C9-HRE-associated diseases have been proposed, including accumulation of nuclear RNA foci containing the C9-HRE RNA, cytoplasmic inclusions of dipeptide repeat-containing (DPR) proteins, and *C9orf72* haploinsufficiency leading to decreased expression of C9orf72 mRNA and protein [8, 9]. In addition, other pathological protein inclusions, such as those containing TAR DNA-binding protein 43 (TDP-43) and p62/SQSTM1, are typically present in the brain of both genetic and sporadic (no clear genetic background) FTD patients [10, 11].

Increasing evidence suggests that structural and functional alterations at synapses and changes in different neurotransmitter systems, such as the glutamatergic, dopaminergic, GABAergic, and serotonergic systems, may underlie the variety of clinical, neuropsychiatric, and cognitive symptoms in FTD and ALS patients [12, 13]. For example, aggression, compulsive behavior, and altered eating habits may arise from disturbances in these systems, suggesting that targeting them might offer new therapeutic options for FTD [12, 14]. Especially, dysfunction in the glutamatergic and GABAergic systems correlate best with disinhibition and behavioral changes in FTD, such as loss of empathy, increasing impulsiveness, and risky decision-making [15, 16]. Both neurophysiological and brain imaging studies have indicated synaptic loss and perturbances in the glutamatergic and GABAergic systems in both sporadic and genetic FTD, including that associated with C9-HRE [17–20]. In C9-HRE-associated FTD, early alterations are observable also in the dopaminergic and cholinergic systems, whereas FTD patients carrying *MAPT* (microtubule associated protein tau) display changes in dopamine and serotonin levels at early stages of disease [21, 22]. Moreover, studies in different cellular and animal models further support the contribution of disturbed glutamatergic and GABAergic neurotransmitter systems in FTD pathogenesis [14, 22–24]. Interestingly, no changes, suggesting alterations in neurotransmitter systems, have been observed in *GRN* (progranulin) mutation carriers in neuroimaging [21]. However, TDP-43 pathology, which is typically present in *GRN* mutation carriers, has been suggested to cause disturbances in the dopaminergic system [16]. Excessive levels of glutamate can lead to synaptic disturbances and trigger excitotoxic neuronal death, involving elimination of GABAergic inhibitory synapses, which subsequently leads to reduced synaptic inhibition and enhanced circuit excitability [25]. Glutamate-mediated excitotoxicity has been observed in C9-HRE-associated FTD, further underlining the involvement of synaptic perturbations in FTD pathogenesis [12].

Synaptic dysfunction is proposed to occur well before symptom onset and to be a unifying early hallmark in FTD and ALS [21]. In early disease stages, ALS patient motor neurons display altered excitability [24, 26]. Similar abnormalities can be detected in FTD-ALS patients and, interestingly, also in FTD patients without clinical evidence of motor symptoms [27, 28]. Moreover, induced pluripotent stem cell (iPSC)-derived neuronal models of C9-HRE have displayed altered synaptic function and neuronal networks [29, 30]. In line with these data, our previous study utilizing overexpression of C9-HRE in mouse primary neurons indicated increased susceptibility of the neurons to glutamate-mediated excitotoxicity, morphological alterations at dendritic spines, and a hyperexcitation phenotype driven by extrasynaptic NMDA receptors [31]. Given all the current information suggesting synaptic dysfunction as an important contributor to FTD pathogenesis, understanding the molecular and cellular mechanisms underlying synaptic changes is crucial for the development of novel therapies aiming at restoring proper synaptic function and improving the clinical outcomes of patients.

Here, our aim was to generate iPSC-derived neurons from both C9-HRE-carrying and sporadic FTD patients and compare their neuropathological and functional features with a special focus on synaptic alterations. We describe both neuropathological as well as structural and functional synaptic alterations in the neurons. The latter were associated with expressional changes in different genes regulating synaptic structures and function as well as altered responses of the neurons to neurotransmitter stimuli, particularly GABA. Moreover, especially the C9-HRE-carrying neurons were found to display nuclear changes in accordance with increased DNA damage. These findings altogether suggest that these patient-derived neurons are excellent models to further explore the molecular mechanisms underlying synaptic disturbances in FTD and that they could potentially be used as platforms for testing different neuromodulatory treatments targeting synaptic dysfunction.

## Materials and methods

### Induced pluripotent stem cell lines and their culture

The iPSC lines used in this study were generated from skin biopsies as described in [32] obtained at the Neuro Center, Neurology, Kuopio University Hospital. All biopsy donors gave written informed consent. The research in human subjects is performed in accordance with the ethical standards of the Declaration of Helsinki and approved by the Research Ethics Committee of the Northern Savo Hospital District, Kuopio, Finland with the ethical permit 16/2013. Studies on FTD patient-derived iPSC-neurons are performed with permission 123/2016 from the Research Ethics Committee of the Northern Savo Hospital District. In this study, iPSCs were generated and characterized from six Finnish FTD patients (53-77 years) clinically diagnosed with bvFTD and three age-matched healthy control individuals according to [33]. The FTD patient iPSCs were derived from three C9-HRE carriers and three sporadic (non-genetic) FTD patients. In addition to those lines, a commercially available line from an FTD patient carrying the C9-HRE was used in some experiments. The line was purchased from EBiSC (cell line name: UCLi001-A, Biosamples ID: SAMEA3174431). EBiSC bank acknowledges University College London and UCL Queen Square Institute of Neurology with support from the NIHR UCLH Biomedical Research Centre, EFPIA companies and the European Union (IMI-JU’). Repeat-primed PCR (AmplideX® PCR/CE C9orf72 Kit, Asuragen) was performed on genomic DNA extracted from corresponding blood samples and iPSCs to confirm the presence or absence of the *C9orf72* HRE in the carriers (C9+,1 line > 60 repeats; 2 lines >145) and sporadic FTD (C9-, < 30 repeats) and healthy controls (< 30 repeats). Patient data were pseudonymized and handled using code numbers. All the iPSC lines were tested negative for mycoplasma regularly (Lonza, LT07-418). iPSCs were maintained on Matrigel (Corning, 356230) coated 3.5 cm dishes (Sarstedt) in E8 medium (Essential 8™ Medium; Gibco,1517001), supplemented with, 1×E8 supplement (Gibco, A1517001) and 0.5% (v/v) penicillin/streptomycin (Pen/Strep; Gibco,15140122) at +37°C and 5% CO_2_. Cell culture medium was replaced every second day. Once confluency of 60-80% was reached, iPSCs were split by incubating for 3 min at +37°C in EDTA (Invitrogen,15575020) containing E8 medium and whole medium was changed on the following day. For freezing, iPSCs were collected in E8 medium supplemented with 10% (v/v) heat inactivated FBS (Gibco,10500) and 10% (v/v) DMSO (Sigma D2650) and stored long-term in liquid nitrogen. After thawing the iPSCs were kept in E8 medium containing 5 µM Y-27632 2HCl (Selleckchem, S1049) for 24h.

### Differentiation of iPSCs to neuronal progenitor cells and cortical neurons

To induce neuronal differentiation, the iPSCs were seeded onto Matrigel-coated 6-well plates in E8 medium. When the colonies reached appropriate size (∼50% confluency), the medium was changed to E8 medium containing the two dual SMAD inhibitors: 10 µM SB431542 (GF-βRI inhibitor, Selleckchem, S1067) and 200 nM LDN193189 2HCl (BMP type I receptor inhibitor, Selleckchem, S7507). The concentration of the dual SMAD inhibitors was kept the same for the whole differentiation process and they were added to the medium every day. On day 1, to start neuronal differentiation, the whole medium was changed containing two parts E8 medium and one part neural differentiation medium (NDM) containing 1:1 DMEM F12 / Neurobasal medium supplemented with 1% B27 (Thermo Fisher, 12587010) 0.5% N2 (Thermo Fisher, 17502048), 2 mM Glutamax (Thermo Fischer, 35050038), and 0.5% (v/v) Pen/Strep. On day two, a full medium change was performed containing two parts NDM medium and one part E8 medium. Starting from day three, the whole medium containing only NDM and the dual SMAD inhibitors was changed. When neuronal rosettes containing differentiated neuroepithelial cells appeared after approximately day 10-12, the medium was changed to NDM containing 25 ng/ml bFGF (100-18B, Peprotech) for about two days, depending on the rosette size and appearance. On day 14, the colonies containing the neuronal rosettes were manually scraped into small clusters and transferred onto ultra-low culture dish (ULA; 4615, Corning) with neuronal sphere medium (NSM) containing 1:1 DMEM F12 / Neurobasal medium supplemented with 0.5% N2, 2 mM Glutamax and 0.5% (v/v) Pen/Strep. On the next day, the whole medium was carefully changed without disturbing the neuronal spheres and from that on, about ¼ of the medium was changed three times a week with NSM supplemented with bFGF. The spheres were manually dissected into smaller spheres to maintain a progenitor-state neural cell population and expanded at least once every 2 weeks. The spheres were cultured for up to eight months. The first experiments were performed after 30 days in culture, but most experiments were performed after 180+ days in culture. The spheres were dissociated with Accutase (Thermo Fischer, A1110501) and plated in NSM medium w/o bFGF onto plated double-coated with 0.1 mg/ml poly-l-ornithine hydrobromide (Sigma, P3655) in Dulbecco’s Phosphate-Buffered Saline (DPBS) overnight at +37°C and 30 µg/ml laminin (Sigma-Aldrich, L2020) in DPBS for four hours (with density 1 × 10^6^ cells/6-well plate or 100,000 cells/24-well plate) to allow acquirement of neuronal morphology. The neurons were maintained for 14 days in NSM medium before performing experiments. Medium samples were collected before fixing the neurons.

### Immunocytochemistry and fluorescence *in situ* hybridization

For immunocytochemistry (ICC), the neurons were fixed with 4% PFA for 20 min at RT and permeabilized in 0.25% Triton X-100 for 60 min at RT and washed twice with DPBS. The cells were blocked in 1% bovine serum albumin (BSA; Sigma) in DPBS for 60 min at RT and incubated with primary antibodies against MAP2 (1:100, Sigma, 9942), GFAP (1:400, Dako, Z0334), vGlut1 (1:300, Sigma, V0389), C9orf72 (1:400, Genetex, GT1553), MAP2 conjugated to CoraLite®594 Fluorescent Dye (1:400,Proteintech, CL594-17490), p62/SQSTM1 (1:200,Santa Cruz, sc-28359), TDP-43 conjugated to CoraLite® Plus 488 Fluorescent Dye (1:100,Proteintech, CL488-80002), or anti-phospho-histone γH2A.X Ser139 (1:400, Sigma, 05-636) overnight at +4°C. The cells were then incubated with fluorescently labelled secondary antibodies Alexa Fluor 647, Alexa Fluor 594, or Alexa Fluor 488 (Invitrogen) for 1 h at RT. Afterwards, the coverslips were washed with DPBS and mounted with Vectashield Vibrance Antifade mounting medium containing 4, 6-diamidino-2-phenylindole (DAPI, Vector Laboratories) and imaged with Zeiss Axio Observer inverted microscope equipped with a Zeiss LSM 800 confocal module (Carl Zeiss Microimaging GmbH, Jena, Germany) or Olympus BX-51 fluorescent microscope with 40X or 60X objective.

For fluorescence in situ hybridization (FISH), diethyl pyrocarbonate DPBS (DEPC-DPBS) and DEPC-treated H_2_O were used. FISH was performed in an RNase-free environment and fluorescently labelled locked nucleic acid (LNA) TYE™ 563-(CCCCGG)_3_ probe (C_4_G_2_; Exiqon) was used to detect the sense foci. TYE™ 563-(CAG)_6_ (CAG; Exiqon) was used as a negative control probe. The cells on coverslips were permeabilized in 0.2% Triton X-100 for 15 min at RT and washed three times in DEPC-DPBS. The coverslips were handled through an ethanol series (70%, 70%, and 100%, 1 min each) followed by drying the coverslips for 10 min at RT. Next, pre-hybridization was performed in hybridization buffer containing 10% dextran sulfate (Millipore, 3730-100ML), 10 mM Ribonucleoside Vanadyl Complex (NEB # S1402S - 200 mM), formamide (Midsci, IB72020), 20 x SSC (Ambion, AM9763), and 1 M sodium phosphate buffer, pH 7, in DEPC-H_2_O, at +66°C for 30 min. The probes were denatured at +85°C for 1 min 15 s and added to hybridization buffer at a concentration of 40 nM C_4_G_2_ or CAG probe and incubated with the coverslips at +55°C for 4 h. Then, after washing in washing buffer 1 (0.1% Tween in 2 X SSC) for 5 min at RT and washing buffer 2 (0.2 X SSC) for 3 x 10 min at +60°C, the coverslips were mounted with Vectashield Vibrance Antifade mounting medium containing DAPI and imaged with Zeiss LSM800 Airyscan confocal microscope.

### Neurofilament light chain Simoa measurements

Neurofilament light chain (NfL) levels in the culture medium were quantified with the Quanterix single molecule array (Simoa, Billerica, MA, USA) HD-X analyzer, using the Simoa® NF-light™ V2 Advantage kit (104073) according to the manufacturer’s instructions. Briefly, frozen medium samples were slowly thawed on ice, mixed, and centrifuged at 10,000 x g for 5 min at RT, and loaded to a 96-well plate. Samples were measured in duplicate and using 1:4 dilution. The Lower Limit of Detection of NfL was reported at 0.038 pg/ml and the Lower Limit of Quantification at 0.174 pg/ml.

### Dipeptide repeat protein measurements

iPSC neurons were cultured for 14 days, then medium was collected and the cells snap frozen. The Mesoscale DPR immunoassay was performed as described in [34]. Briefly, the frozen cell pellets were thawed on ice for 10 min and lysed in RIPA buffer. DPR immune assay was performed on the MSD platform using streptavidin plates (MSD Gold 96-well Streptavidin SECTOR: #L15SA). The plates were coated overnight with biotinylated capture antibody in blocking buffer (PBS with 0.05% Tween-20, 1% BSA) at +4°C. After washing (DPBS with 0.05% Tween-20), the plates were incubated for 2 h at RT on a shaking platform (140 rpm). After washing, sulfo-tag-labeled detection antibody diluted in blocking buffer was added and the plates were incubated for 2 h at RT on a shaking platform (140 rpm), followed by washing. The electrochemiluminescent signal was measured using a MESO QuickPlex SG120 instrument. Antibodies used for poly-GP measurement were biotinylated 18H8 (0.13 mg/ml) and sulfo-tagged 3F9 (1 mg/ml). Antibodies for poly-GA measurement were biotinylated 1A12 (1 mg/ml) and sulfo-tagged 1A12 (1 mg/ml). Total protein levels in all lysates were quantified using PierceTM BCA Protein Assay Reagents (Thermo Fisher: #23222 and #23224). After the incubation step for the poly-GP immunoassay, a fraction of the lysed sample was used to measure total protein concentration. The DPR antibodies were a kind gift from Prof. Dr. Dieter Edbauer.

### 3D culture and dendritic spine analysis

For dendritic spine analysis, the 10-mm microwell in a 3.5-cm glass-bottom microwell dishes (MatTek, P35G-1,5-10-C) were coated with 0.1 mg/ml poly-l-ornithine hydrobromide for two hours at +37°C. Then, the coating was removed, and the dish left to dry overnight in the in the laminar at RT. Neuronal spheres were collected and dissociated using Accutase (∼500,000 cells/0.5ml) and mixed with green fluorescent protein (GFP) adeno-associated virus with multiplicity of infection (MOI) 5,000. The GFP construct was packaged in AAV9 virus particles at the BioCenter Finland National Virus Vector Laboratory, University of Eastern Finland, Kuopio, Finland. Matrigel and cell/virus suspension was mixed in a ratio of 1:8 and a 100-µl drop was added to the coated microwells and incubated at +37°C and 5% CO_2_ for at least 24 h to harden the Matrigel. Then NSM medium was carefully added. Half of the medium was changed three times per week. The cultures were cultured for additional up 180 days for dendritic spine analysis. Dendritic spines from GFP-positive neurons were imaged with LSM 800 Airyscan confocal microscope. Serial Z-stacks of optical sections from dendritic segments were analyzed by NeuronStudio software (Computational Neurobiology and Imaging Center Mount Sinai School of Medicine, New York, Version 0.9.92 64-bit). Settings were adjusted as following: Volume: pixel dimensions were set to X = 0.066 μm (pixel width), Y = 0.066 μm (pixel height), and Z = 0.390 μm (voxel depth); Dendrite detection: attach ratio was set to 1.5, minimum length to 3 μm, discretization ratio to 1, and realign junctions to yes; Spine detection: maximum height of spines was set to 5.0 μm and minimum height to 0.2 μm, maximum width to 3 μm, maximum stubby size to 10 voxels, and minimum non-stubby size to 5 voxels; Spine classifier: neck ratio (headneck ratio) was set 1.1, thin ratio to 2.5, and mushroom size to 0.35 μm. To obtain a clear image, the image filter Blur-MP was run before the analysis and the dendritic spines were sub-grouped according to their morphology to mushroom, stubby, or thin spines according to Neuron studio’s automatic settings using the Rayburst algorithm.

### Calcium imaging

iPSC neurons were loaded with calcium-sensitive fluorescent Fluo-4 dye (1×, Direct Calcium Assay Kit, Invitrogen, USA) in humidified incubator (5% CO2, +37°C) for 30 - 40 min following by washout for 10 min at +37°C and then for 5 min at RT in basic solution (BS) containing 152 mM NaCl, 2.5 mM KCl, 10 mM HEPES, 10 mM glucose, 2 mM CaCl2, 1 mM MgCl2, pH adjusted to 7.4. Then, coverslips with the loaded cells were transferred to the recording chamber with constant BS perfusion (flow rate 2.5 ml/min, RT). Fluo-4 loaded cells were imaged using monochromatic light source (TILL Photonics GmbH) with an excitation light wavelength of 495 nm (exposure time 100 ms) and registered emission intensity at ≥520 nm. Fluorescence was visualized with 10× objective in Olympus IX-70 microscope equipped with CCD camera (SensiCam, PCO imaging, Kelheim, Germany) and recorded with TILL Photonics Live Acquisition at 1-Hz sampling frequency (1 frame per second). Setup was equipped with Rapid Solution Changer RSC-200 (BioLogic Science Instruments, Grenoble, France) allowing local application of various solutions, with fast exchange between them (∼30 ms). The cells were sequentially treated with short (2 s) applications (separated by 2 min BS washout) of 100 μM GABA (Sigma), 100 μM glutamate (together with 30 μM NMDA receptor co-agonist glycine; both from Sigma), 50 mM KCl (depolarizing agent, to differentiate neuronal cells from glia), and 10 μM ionomycin (membrane-permeable calcium ionophore, to define the cells’ viability; Tocris).

### Analyses of C9orf72 protein levels, TDP-43 nucleus-to-cytoplasmic translocation, and p62/SQSTM1 vesicles

iPSC neurons were stained with antibodies against MAP2, C9orf72, and with DAPI, as described above. C9orf72 images were acquired using Zeiss LSM800 Airyscan confocal microscope. For the C9orf72 analysis, binary MAP2 masks were created by applying the Li thresholding method to the MAP2 images. Particles smaller than 2 µm were removed and holes within the cell bodies of a size between 0-4 µm^2^ were filled. To capture C9orf72 signal from the full cytoplasm, nuclear masking was applied. DAPI images were used to create nuclear masks by using the Otsu thresholding method, followed by removal of particles with a radius smaller than 5 µm. The C9orf72 signal was quantified as sum intensities of the fluorescence signal originating from the fluorescently labeled secondary antibody within MAP2-positive and DAPI-positive areas (defining the neuronal boundaries) per image. Sum intensities were normalized to MAP2-positive area per image. Images were processed using Fiji (2.3.0/1.53f51; [35]). Neurons were stained with MAP2 and TDP-43 or p62/SQSTM1 antibodies and DAPI, as described above. TDP-43 images were acquired using Zeiss LSM800 Airyscan confocal microscope. p62/SQSTM1 images were acquired using Olympus BX-51 fluorescent microscope. Images were processed using Fiji (2.3.0/1.53f51; [35]). For both analyses, MAP2 images were used to outline neurons and measure the total cell body area per image of neurons only. For TDP-43 analysis, MAP2 images were processed by enhancing the local contrast (blocksize=127, histogram=256, maximum=3.10). Next, Otsu thresholding method was applied, followed by removal of particles with a radius smaller than 5 µm. To fill holes within cell bodies, particles of a size between 0-4 µm^2^ were included. After watershed segmentation, regions of interest were outlined without further area or shape exclusion. Sum intensity values of depicted area from TDP-43 images were acquired. Then, DAPI images were analyzed including only the nuclei of MAP2-positive cells. DAPI images were used to indicate nuclei and measure nuclear areas per image (for TDP-43 analysis). To segment nuclei, thresholding (Renyi’s entropy method) was applied, followed by the fill holes command. Finally, sum intensity values of DAPI-positive areas were extracted from TDP-43 images. TDP-43 signals were quantified as sum intensities of the fluorescence signal originating from the fluorescently labeled antibody targeted against TDP-43 within nuclear (DAPI-positive) and cytosolic areas (nuclear signal subtracted from signal within whole cell body [MAP2-positive areas]) respectively for each image. Sum intensities were normalized to nuclear and cytosolic area sizes respectively for each image. Cytosolic to nuclear TDP-43 signal ratio was calculated for each image of each cell line.

For p62/SQSTM1 image analysis, MAP2 images were converted into binary images using the Li thresholding method, and further processed by removing small objects (≤ 5 µm radius) and using watershed segmentation. Next, p62/SQSTM1-positive vesicles were analyzed from MAP2-positive regions by converting images into binary images using thresholding (Renyi’s entropy method). Only particles with a minimum area of 0.01 µm^2^ to 1 µm^2^ in size were included in the analysis. Number of p62/SQSTM1-positive vesicles per image was normalized to total MAP2 area of the same image. Average size and integrated density of p62/SQSTM1-positive vesicles per image and cell line were extracted from Fiji.

### Nuclear morphology and DNA damage analysis

iPSC neurons were stained with γH2A.X antibody and DAPI as described above. Nuclear masking used in γH2A.X analysis was created in Fiji based on the DAPI staining by applying unsharp mask (radius = 1, mask = 0.60), followed by enhancing local contrast (CLAHE) (blocksize = 127, histogram = 256, maximum = 3) and adding a median filter (radius = 3). After converting the image to binary, particles with a radius smaller than 5 µm were removed. Total nuclear area per image was acquired by measuring masked regions with an area of 1.5 µm^2^ or higher. For γH2A.X masking, any signal in the γH2A.X channel outside of the DAPI mask was removed. The remaining signal was measured as sum intensity of the fluorescence signal from the fluorescently labeled secondary antibody, and the sum intensity from one image was normalized to the total nuclear area in the image. To acquire γH2A.X-positive particle number and size, the remaining γH2A.X signal was transformed into binary by auto thresholding (MaxEntropy algorithm), followed by analyzing the particles with an area higher than 0.00001 µm^2^. Particle number per image was normalized to the total nuclear area in the image, and particle size was measured as the mean particle area (µm^2^) per image.

For the analysis of nuclear morphology, the neurons were stained with DAPI as described above and imaged with Zeiss LSM800 Airyscan confocal microscope. To segment the nuclei, the Fiji plugin StarDist [36] was utilized. From StarDist, the Versatile (fluorescent nuclei) built-in model was used, with the non-maximum suppression postprocessing parameters of 0.5 (Probability/Score threshold) and 0.6 (Overlap threshold). The acquired nuclear regions-of-interest were copied to the non-segmented image, and morphological parameters of each DAPI-positive particle were measured. The following morphological parameter was calculated per DAPI-positive particle: eccentricity [sqrt(a^2^-b^2^)/a], where a = major axis length and b = minor axis length.

The images were divided into two most similar groups based on the mean area of the DAPI-positive particles per image using K-means clustering. Python version 3.11.8 and the following libraries were used for clustering. The data from each group was reshaped with the NumPy (version 1.26.2) module “reshape” using numpy.reshape (−1,1) prior to clustering. Data points from each group separately were clustered into two groups using the KMeans algorithm from the scikit-learn [27][45], (version 1.4.1) module “sklearn.cluster”.

To study the DNA damage response, iPSC-derived neurons were treated with 10 µl topotecan (Topotecan hydrochloride, Tocris, 4562) or DMSO vehicle for 1h. The cells were fixed with 4% PFA and stained with conjugated MAP2 and γH2A.X antibody as described above. Images were processed using Fiji (2.3.0/1.53f51). The nuclear masking used for γH2A.X analysis was based on the DAPI staining. It was created by applying unsharp mask (radius = 1, mask = 0.60), followed by enhancing local contrast (CLAHE) (blocksize = 127, histogram = 256, maximum = 3) and a median filter (radius = 3). The image was converted to binary and particles with a radius smaller than 5 µm were removed. The total nuclear area per image was quantified by measuring masked regions with an area of 1.5 µm^2^ or higher. To mask the γH2A.X, only signals within the DAPI area were used. The remaining signal was measured as sum intensity of the fluorescence signal from the fluorescently labeled secondary antibody, and the sum intensity from one image was normalized to the total nuclear area in the image. To acquire γH2A.X-positive particle number and size, the remaining γH2A.X signal was transformed into binary by auto thresholding (MaxEntropy algorithm), followed by analyzing the particles with an area higher than 0.00001 µm^2^. Particle number per image was normalized to the total nuclear area in the image, and particle size was measured as the mean particle area (µm^2^) per image.

### RNA sequencing and data processing

RNA was extracted using the Direct-zol RNA Miniprep Plus (Zymo, R2072, USA, California). Bulk RNA sequencing (RNA-seq) was performed using RNA extracts from iPSCs and iPSC neurons. Library preparation and RNA sequencing was conducted by Novogene (UK) Company Limited. In brief, mRNA enrichment was performed with oligo(dT) bead pulldown, from where pulldown material was subjected to fragmentation, followed by reverse transcription, second strand synthesis, A-tailing, and sequencing adaptor ligation. The final amplified and size selected library comprised of 250-300 base pair (bp) insert cDNA and paired-end 150 bp sequencing was executed with an Illumina high-throughput sequencing platform. Sequencing yielded 22.3–27.1 million sequenced fragments per sample.

The 150-nucleotide pair-end RNA-seq reads were quality-controlled using FastQC (version 0.11.7) (https://www.bioinformatics.babraham.ac.uk/projects/fastqc/). Reads were then trimmed with Trimmomatic (version 0.39) [38] to remove Illumina sequencing adapters and poor quality read ends, using as essential settings: ILLUMINACLIP:2:30:10:2:true, SLIDINGWINDOW:4:10, LEADING:3, TRAILING:3, MINLEN:50. Reads aligning to mitochondrial DNA or ribosomal RNA, phiX174 genome, or composed of a single nucleotide, were removed using STAR (version 2.7.9a) [39]. The remaining reads were aligned to the Gencode human transcriptome version 38 (for genome version hg38) using STAR (version 2.7.9a) [39] with essential non-default settings: --seedSearchStartLmax 12, --alignSJoverhangMin 15, --outFilterMultimapNmax 100, --outFilterMismatchNmax 33, -- outFilterMatchNminOverLread 0, --outFilterScoreMinOverLread 0.3, and --outFilterType BySJout. The unstranded, uniquely mapping, gene-wise counts for primary alignments produced by STAR were collected in R (version 4.1.0) using Rsubread::featureCounts (version 2.8.2) [40], ranging from 17.2 to 20.7 million per sample. Differentially expressed genes (DEGs) between experimental groups were identified in R (version 4.2.0) using DESeq2 (version 1.36.0) [41] by employing Wald statistic and lfcShrink for FC shrinkage (type=“apeglm”) [42], and correcting for sequencing batch. Pathway enrichment analysis was performed on the gene lists ranked by the pairwise DEG test log2FCs in R using clusterProfiler::GSEA (version 4.4.4) [43] with Molecular Signatures Database gene sets (MSigDB, version 2022.1) [44].

### Data processing, visualization, and statistical analyses

To test whether data points within the experimental groups were normally distributed, Shapiro-Wilk test was used. To test for significance between two different experimental groups, two-tailed independent samples t-test was used for normally distributed data and Mann Whitney U test for not normally distributed data. To test for significance between more than two groups, one-way analysis of variance (ANOVA) followed by Tukey’s multiple comparisons test or Sidak’s multiple comparison test was used if data points were normally distributed. Otherwise, Kruskal-Wallis test followed by Dunn’s multiple comparisons test was used. For grouped variables, two-way analysis of variance (ANOVA) followed by Tukey’s multiple comparisons test was used. To assess statistical independence between categorical variables, the Chi-Square test was used. All tests were performed using GraphPad Prism software (version 8.4.3 for Windows, GraphPad Software, San Diego, California USA, www.graphpad.com). To check for and remove potential outliers in the data, GraphPad Prism’s Outlier Identification (ROUT method, Q = 1%) was used. The data are shown as mean ± standard deviation (SD) or median ± interquartile range.

For calcium imaging, the data were post-processed using FEI offline analysis (TILL Photonics GmbH, Germany) and pre-analyzed offline with Image J (Rasband, W.S., Image J, U.S. National Institutes of Health, Bethesda, Maryland, USA). Final analysis and plotting were performed using Origin 2019b software (OriginLab Corporation, Northampton, Massachusetts, USA). To quantify the amplitude of calcium responses, the ratio of the transient fluorescence response to baseline fluorescence was determined (ΔF/F0, the signal-to-baseline ratio; ΔF/F0 = (F-F0)/F0, where F is the calcium-transient peak, and F0 is the averaged baseline fluorescence under resting conditions). Quantitative data are expressed as means ± standard error of mean (SEM), unless otherwise stated. Number of batches and cells within each batch were indicated by n. Significance was assessed with Student’s paired t-test and one-way ANOVA followed by Tukey post hoc test for parametric and non-parametric data, respectively.

P-values ≤ 0.05 were considered significant. Statistically significant differences are shown as *p < 0.05, **p <0.01, and ***p <0.001. Graphs were created using the GraphPad Prism software (version 8.4.3 for Windows. Total number of statistical units per group is indicated as “n”. Statistical units are depicted as individual data points.

## Results

In this study, we differentiated iPSCs from healthy donors (controls) and sporadic as well as C9-HRE-carrying FTD patients to cortical neurons using the dual SMAD inhibitors. Synaptic dysfunction has been implicated in FTD pathogenesis, but it has remained unclear whether the neurons and synapses are similarly affected in all FTD patients or whether there are specific alterations depending on the genetic background of the patient. Here, we report our findings related to the cell pathological phenotypes and synaptic alterations at the structural, functional, and gene expression level. Prompted by the global RNA-seq data, we also compared the potential nuclear alterations between the sporadic and C9-HRE FTD and control neurons.

### Characterization of the iPSC-derived cortical neuron cultures

First, we characterized the obtained iPSC-derived neuronal cultures. Morphologically, there were no obvious differences between the neuronal cultures from sporadic and C9-HRE-carrying FTD patient neurons and those from the healthy control individuals (**Figure 1A**). ICC staining indicated that the cultures after 180 days contained glutamatergic and GABAergic MAP2-positive neurons as well as GFAP-positive astrocytes. We also evaluated the success of the neuronal differentiation process by comparing gene expression of specific marker genes between iPSCs and iPSC-derived neurons from the global RNA-seq data (**Figure 1B**). Stem cell markers, such as NANOG and POU5F1, which play a key role in embryonic development and stem cell pluripotency, were downregulated in the neurons as compared to iPSCs. The expression levels of SOX2, a key factor regulating pluripotency and neural differentiation [45], were similar in iPSCs and the neurons. PAX6, a marker for both stem cells and neuroectodermal differentiation has previously been reported to be upregulated after dual SMAD inhibition [46, 47]. Here, it was also expressed at higher levels in the differentiated neurons than in the iPSCs. The neuronal marker MAP2, essential for neurogenesis, was upregulated in the iPSC-derived neurons as compared to the iPSCs. In addition, the neuronal cultures expressed markers from all six cortical layers of the human brain and these markers were low in the iPSCs (**Figure 1B**). BLC11B represents neurons from layers 5-6, SATB2 marks a distinctive subpopulation of upper layer neurons, POU3F2 is found in layer 2/3, 4 and 5 neurons, TUBB3 is related to glutamatergic synapse function, EOMES supports cortical neurogenesis expansion, and TBR1 is found in layer 6 cortical neurons. [48–52]. In addition, astrocyte markers GFAP and S100B showed upregulated levels after 180 days in culture (**Figure 1B**). Measurement of released NfL, a generally clinically used biomarker of neurodegeneration, in the conditioned media of the neurons using Simoa indicated undetectable levels of NfL, suggesting that there was no observable neuronal cell death in any of the cultures (data not shown).

**Figure 1:**
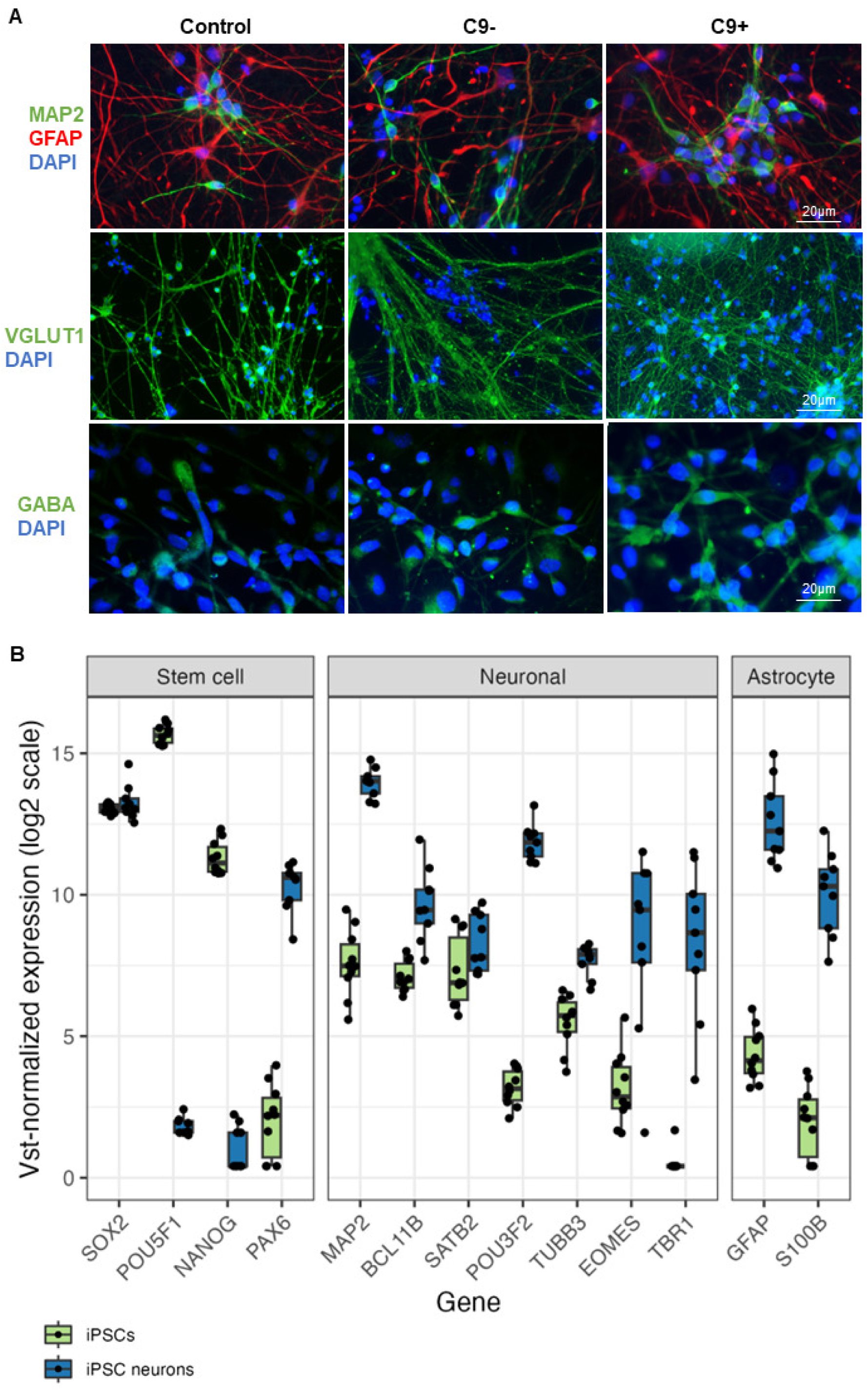
Characterization of the iPSC-derived neurons derived from sporadic and C9-HRE-carrying FTD patients and healthy control individuals. Immunocytochemical staining of the iPSC neurons indicated that the cultures after 180 days encompass MAP2-positive neurons (green) and GFAP-positive astrocytes (red) (**A**). Also, the neurons are positive for vGLUT1 and GABA staining, indicating the presence of glutamatergic and GABAergic neurons, respectively (A). Representative microscopy images are shown. RNA sequencing confirmed successful differentiation of the iPSCs to cortical neurons (**B**). Stem cell markers NANOG and POU5F1 decreased in the iPSC neurons. Neuronal marker expression, such as that of MAP2 and PAX6, a marker for both stem cells and neuroectodermal differentiation, increased in the iPSC-derived neurons. The neuronal cultures expressed markers from all six cortical layers of the human brain (BLC11B, SATB2, POU3F2, TUBB3, EOMES and TBR1) and these markers remained low in the iPSCs. Astrocyte markers GFAP and S100B also increased in the cultures after 180 days.

### The C9-HRE-carrying FTD iPSC neurons exhibit C9-HRE-associated pathological hallmarks

Next, we studied if the C9-HRE FTD patient derived neurons presented the typical C9-HRE-associated gain and loss-of-function pathological hallmarks. FISH analysis using the TYE™ 563- (CCCCGG)_3_ probe, recognizing the sense strand-derived expanded GGGGCC repeat-containing RNA, revealed that only the C9-HRE-carrying but not the control or sporadic FTD neurons displayed nuclear RNA foci (**Figure 2A**). Importantly, no RNA foci were detected in C9-HRE neurons using the TYE™ 563-(CAG)_6_ negative control probe, indicating specificity of the TYE™ 563-(CCCCGG)_3_ probe against the expanded GGGGCC repeats (data not shown). Mesoscale-based DPR immunoassay revealed that the poly-GP (**Figure 2B**) and poly-GA (**Figure 2C**) DPR proteins were specifically detected in all C9-HRE-carrying neurons. Quantitative image analysis of C9orf72 ICC indicated that the sporadic FTD neurons had the highest levels of C9orf72 protein, and these were significantly higher than those in the control or C9-HRE FTD neurons. There were no statistically significant differences in the C9orf72 protein levels between the C9-HRE FTD and control neurons, suggesting that the C9-HRE neurons do not show signs of *C9orf72* haploinsufficiency, although the levels appeared the lowest in the C9-HRE-carrying neurons (**Figure 2D**).

**Figure 2:**
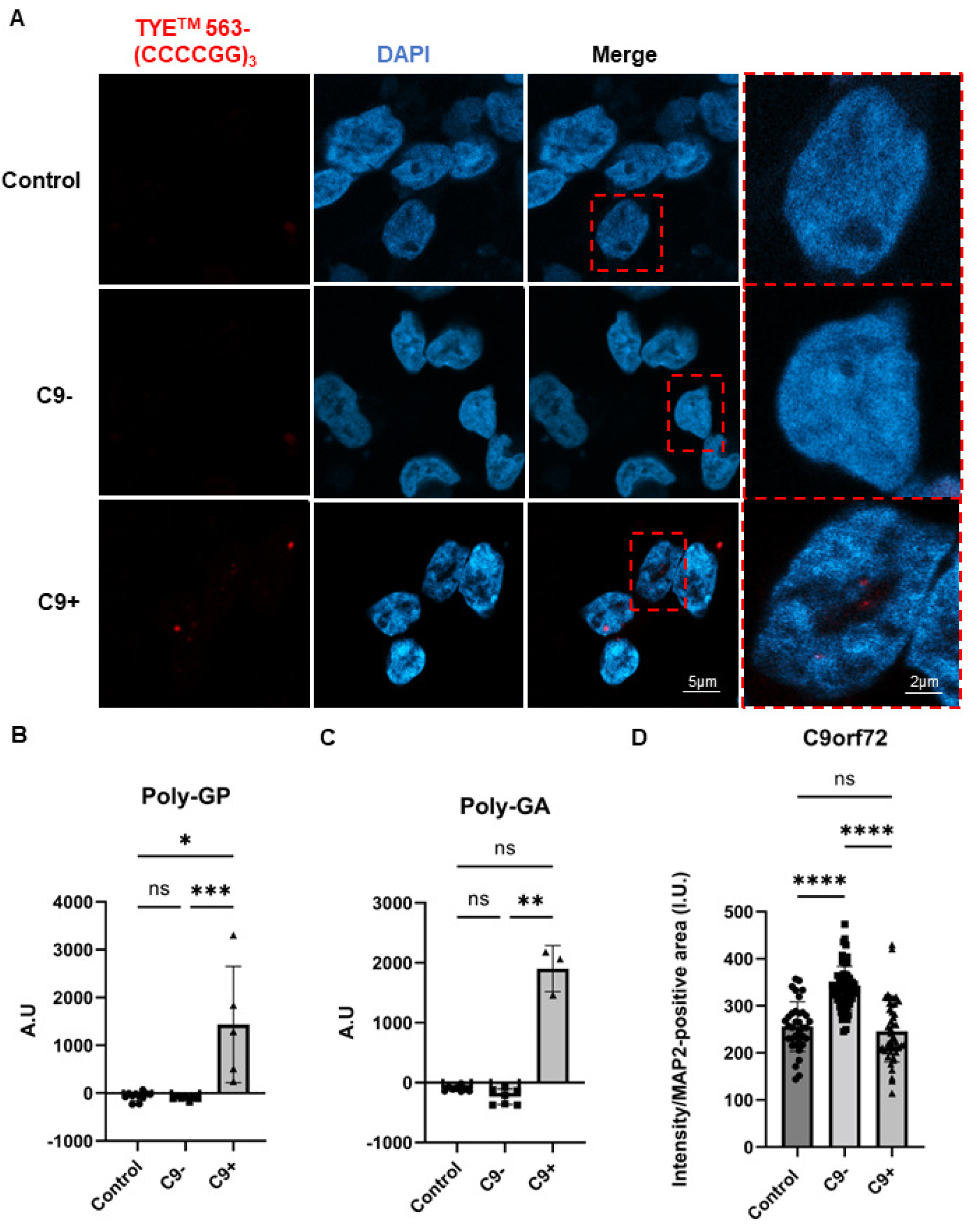
iPSC-derived neurons from C9-HRE-carrying FTD patients express typical C9-HRE-associated pathological hallmarks. Fluorescence *in situ* hybridization (FISH) shows that C9-HRE neurons but not sporadic FTD or control neurons exhibit sense RNA foci (red) as detected using the TYE^TM^ 563-(CCCCGG)_3_ probe (**A**). Representative microscopy images are shown. Also, C9-HRE carrying neurons specifically produce poly-GP (**B**) and poly-GA (**C**) DPR proteins. One-way ANOVA followed by Tukey’s multiple comparisons test was used. *n* [Control] = 11; *n* [C9-] =8; *n* [C9+] = 5. C9-HRE-carrying neurons do not present with C9orf72 protein haploinsufficiency but a slight non-significant trend towards lower C9orf72 protein levels as compared to controls (**D**). One-way ANOVA followed by Tukey’s multiple comparisons test was used. *n* [Control] = 35; *n* [C9-] = 61; *n* [C9+] = 46. Statistically significant differences are shown as *p < 0.05, **p <0.01, and ***p <0.001.

### iPSC-derived FTD neurons display altered intracellular TDP-43 localization and enlarged p62/SQSTM1-positive vesicles

TDP-43 pathology indicated by the accumulation of TDP-43 in cytoplasmic inclusions is a major hallmark of FTD brain, especially in FTD associated with the C9-HRE. ICC revealed that the FTD patient-derived neurons did not display cytosolic TDP-43 inclusions (**Figure 3A**). However, quantitative image analysis revealed that in the FTD neurons, regardless of their C9-HRE carriership status, the cytosolic-to-nuclear ratio of TDP-43 was significantly increased compared to the control neurons (**Figure 3B**), suggesting increased nucleus-to-cytosolic shuttling of TDP-43. Moreover, we observed that the size of the p62/SQSTM1-positive vesicles was significantly enlarged and their intensity was stronger in both sporadic and C9-HRE-carrying FTD neurons compared to controls (**Figure 3C-E**). However, the FTD neurons displayed a trend towards a lower number of p62/SQSTM1-positive vesicles compared to controls, but this remained statistically non-significant (Figure 3F). These results suggest that in the FTD neurons, there are fewer, but larger vesicles containing increased levels of p62/SQSTM1 compared to neurons of healthy controls.

**Figure 3:**
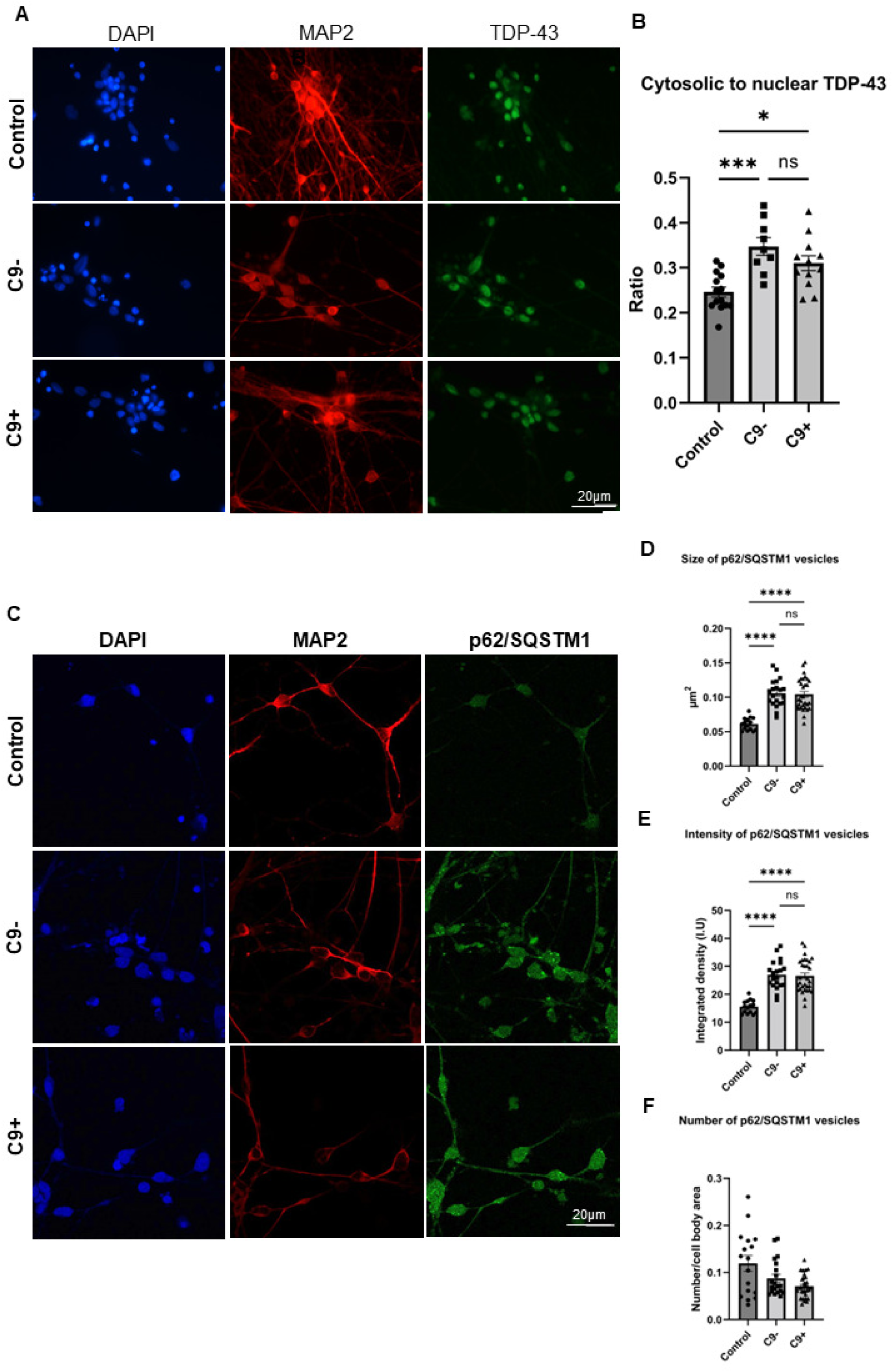
iPSC-derived neurons from C9-HRE-carrying as well as sporadic FTD patients show altered intracellular TDP-43 shuttling and enlarged p62/SQSTM1-positive vesicles. Representative microscopy images of TDP-43-stained (green) neuronal cultures are shown (**A**). Quantification of cytosolic-to-nuclear ratio of TDP-43 based on immunofluorescence imaging (**B**). Each data point represents cytosolic-to-nuclear TDP-43 signal from one image. For each group, descriptive statistics are shown as mean ± standard deviation. One-way ANOVA followed by Tukey’s multiple comparisons test was used. *n* [Control] = 14; *n* [C9-] = 9; *n* [C9+] = 12. DAPI (blue) and MAP2 (red) signals were used to outline neuronal nuclei and cell boundaries, respectively, and to distinguish the TDP-43 signal in the nucleus and in the cytosol. Quantification of p62/SQSTM1 based on immunofluorescence imaging. Each data point represents size of p62/SQSTM1 vesicles (**D**), fluorescence intensity of p62/SQSTM1 vesicles (**E**), and p62/SQSTM1-positive vesicles per MAP2-positive area per image (**F**). For each group, descriptive statistics are shown as median ± interquartile range. Kruskal-Wallis test followed by Dunn’s multiple comparisons test was used. *n* [Control] = 17; *n* [C9-] = 17; *n* [C9+] = 21. Representative microscopy images of p62/SQSTM1-stained neuronal cultures are shown (**C**). MAP2 signal was used to outline the neuronal cell boundaries, and to normalize the number of p62/SQSTM1-positive vesicles to neuronal cell area. Statistically significant differences are shown as *p < 0.05, **p <0.01, and ***p <0.001.

### iPSC-derived FTD neurons show altered dendritic spine number and morphology as well as decreased response to GABA stimulus

Our previous studies in mouse primary neurons overexpressing C9-HRE showed altered dendritic spine morphologies [31]. Thus, we next explored whether morphological changes in the dendritic spines of the FTD patient-derived neurons can be observed by transducing neurons with GFP-labeled AAV9 virus (**Figure 4A**). Quantitative analysis of the number and morphology of the dendritic spines revealed that the total spine density was significantly reduced in both sporadic and C9-HRE-carrying FTD neurons compared to control neurons (**Figure 4B**). In addition, both sporadic and C9-HRE FTD neurons exhibited a significant decrease in the number of mushroom-type spines compared to control neurons (**Figure 4C**), while the stubby spine density was significantly increased only in the C9-HRE-carrying FTD neurons (**Figure 4E**). The density of thin spines was not affected in the FTD neurons (**Figure 4D**). Interestingly, the head diameter of the mushroom and thin spines was not affected, implicating that the remaining spines may function normally (**Figure 4F-H**). iPSC-derived neurons from sporadic FTD patients displayed a slightly smaller head diameter of the stubby spines (**Figure 4H**). Altogether, these results indicate loss of dendritic spines and significant alterations in the dendritic spine morphologies in iPSC-derived neurons from FTD patients and suggest that alterations in the structure and function of synapses is a common feature in FTD patient-derived neurons.

**Figure 4:**
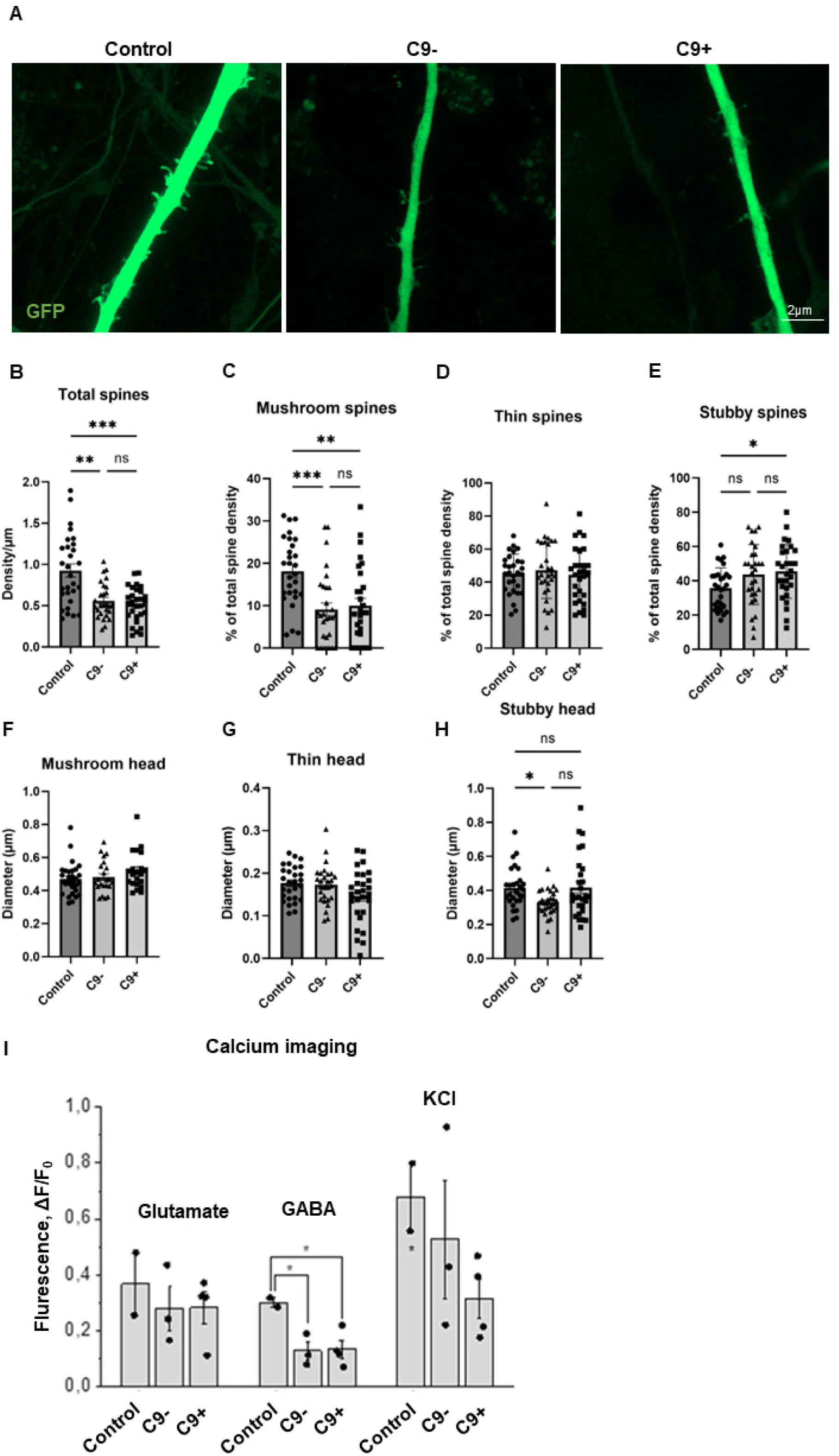
iPSC-derived FTD neurons show altered dendritic spine number and morphology as well as a decreased response to GABA stimulus. Representative images of dendrites and their spines of control, sporadic and C9-HRE neurons are shown (**A**). The neurons expressing GFP (green) and imaged with Zeiss LSM 800 confocal microscope and analyzed with NeuronStudio program. Dendritic spine analysis shows that the total spine number is significantly reduced in iPSC-derived FTD neurons compared to control neurons (**B**). The FTD neurons exhibit significantly fewer mushroom-type spines, but the number of other spine types remains the same, except for the number of stubby spines, which increased only in C9-HRE carriers (**C-E**). The number of each spine type was normalized to total spine number. The head diameter is not altered in any dendritic spine type (**F-G**), except for the sporadic FTD neurons, which display a slight decrease in stubby spine head diameter (**H**). One-way ANOVA followed by Tukey’s multiple comparisons test was used or for not normally distributed data Kruskal-Wallis test followed by Dunn’s multiple comparisons test was used *n* [Control] = 28-30; *n* [C9-] = 21-29; *n* [C9+] = 23-29. Calcium imaging revealed that both sporadic and C9-HRE-carrying FTD neurons display a significantly lower response to GABA compared to control neurons (**I**). Also, the FTD neurons show a non-significant trend to a weaker response to glutamate and depolarizing KCl as compared to control neurons. Quantitative data are expressed as means ± SEM (standard error of mean), unless otherwise stated. Number of batches and cells within each batch were indicated by n. Significance was assessed with Student’s paired t-test and one-way ANOVA followed by Tukey post hoc test for parametric and non-parametric data, respectively. *n* [Control] = 2 (2 lines included per batch); *n* [C9-] = 3 (3 lines included per batch); *n* [C9+] = 3 (3-4 lines included per batch), n indicated averaged numbers from different batch measurements. Statistically significant differences are shown as *p < 0.05, **p <0.01, and ***p <0.001.

Loss of dendritic spines might indicate disturbed synaptic function. Therefore, we next investigated if the structural changes and reduced spine density were associated with altered neuronal functionality. We performed calcium imaging in the iPSC-derived neurons and assessed their responses to short application of glutamate, GABA, and KCl. Both sporadic and C9-HRE-carrying FTD neurons displayed a significantly lower response to GABA compared to control neurons (**Figure 4I**). Also, the FTD neurons showed a weaker response to glutamate and reacted less strongly to depolarizing KCl as compared to control neurons, but these changes did not reach statistical significance.

### Global RNA sequencing reveals several altered biological pathways in iPSC-derived FTD neurons as compared to control neurons but only one differentially expressed gene between sporadic and C9-HRE FTD neurons

Global RNA-seq was performed to study the potential gene expression changes in the FTD patient neurons. The results indicated that the C9-HRE and sporadic FTD neurons displayed largely similar gene expression patterns (**Figure 5A**). However, the gene expression signature of the FTD neurons was dramatically different from that in the control neurons. Several pathways were significantly downregulated in the FTD neurons when compared to control neurons. A more detailed pathway analysis indicated prominent changes in pathways related to synaptic function and signaling (upregulation) as well as DNA repair mechanisms (downregulation) in FTD neurons when compared to control neurons (**Figure 5B**). The most significant changes in upregulated DEGs were observed in genes associated with synaptic function, such as GABA receptors 1 and 2 (*GABRG1* and *GABRG2*), voltage-gated potassium channel subfamilies (*KCND*, *KCNN3*, *KCNK2*), AMPA receptor subunit (*GRIA2*), NMDA receptor subunit (*GRIN2B*), cell-surface receptor neurexin 1 (*NRXN1*), and genes related to neural growth, differentiation, cell adhesion, and signal transduction (*CSPG5* and *CD38*). Significantly downregulated DEGs in the DNA repair mechanism pathway included methyltransferases involved in transfer of methyl groups (*PRMT5*), pro-apoptotic genes, such as (*PMAIP1*), response to DNA damage (*CHEK2*), and genes encoding mitochondrial proteins (*PRELID1*).

**Figure 5:**
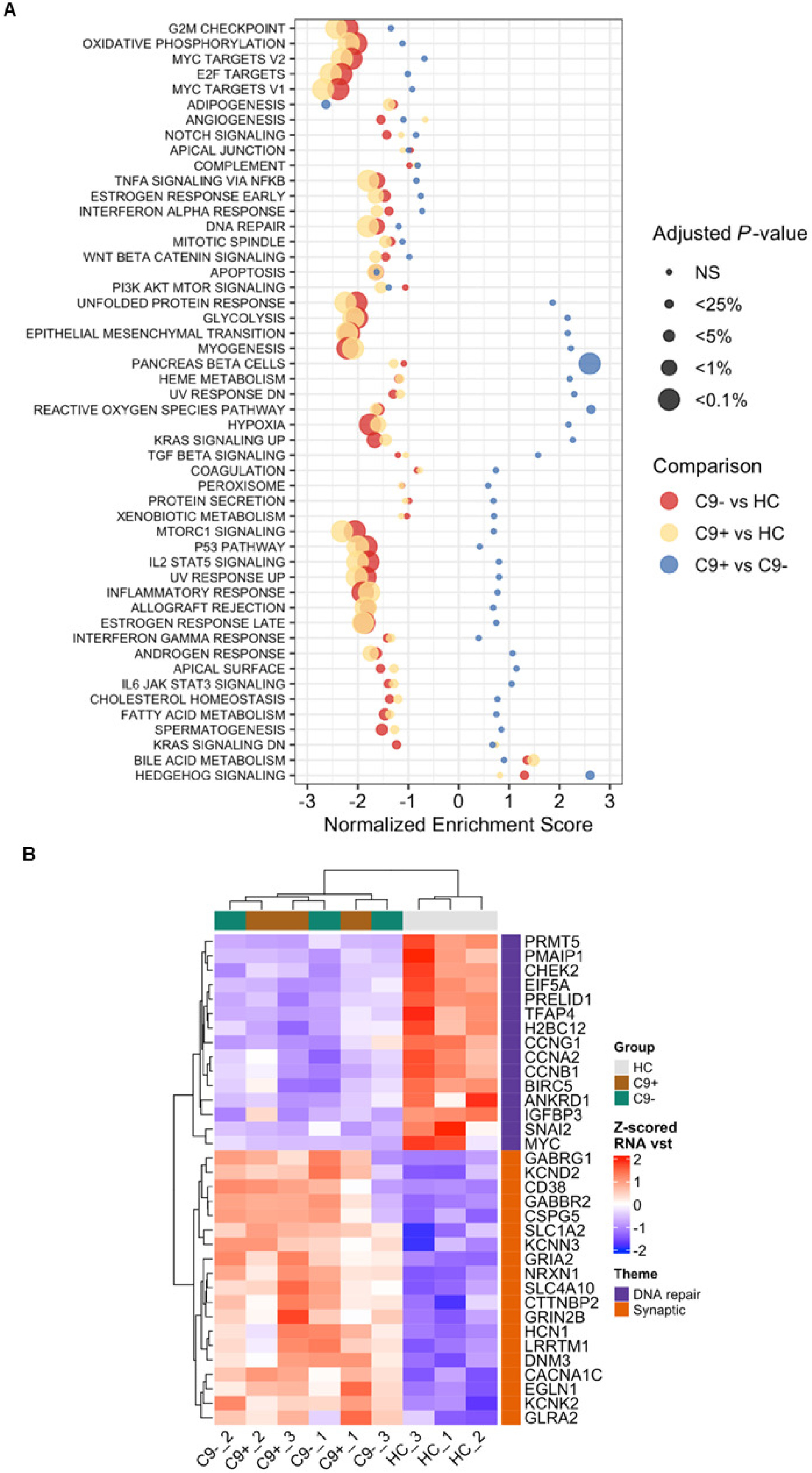
Global RNA sequencing reveals several altered biological pathways in iPSC-derived FTD neurons as compared to control neurons. Global RNA sequencing indicates that the C9-HRE and sporadic FTD neurons display largely similar gene expression patterns (**A**). However, the gene expression signatures of both the sporadic and C9-HRE-carrying FTD neurons are dramatically different from those in the control neurons. The dot plot displays, for the three differently colored comparisons, the gene set enrichment analysis (GSEA) results for the MSigDB gene set collections GO Biological Processes (C5 GO BP) and Canonical Pathways (C2 CP). Normalized enrichment score (NES) is shown in the x-axis whereas the categorical statistical significance (FDR) is indicated by the size of the marker. Several pathways are significantly downregulated in the FTD neurons when compared to control neurons. A more detailed pathway analysis indicated prominent changes in pathways related to synaptic function and signaling (upregulation) as well as DNA repair mechanisms (downregulation) in FTD neurons when compared to control neurons (**B**). Heatmap shows the row-wise Z-scored vst-normalized expression in individual samples for genes that were highly significant up or down-regulated C9+ vs. HC DEGs (P-adj. < 0.01) and among the GSEA core enrichment of either select synaptic (for up-regulated DEGs) or DNA repair related (for down-regulated DEGs) GO or Canonical pathways.

When comparing the C9-HRE and sporadic FTD neurons, it was interesting to note that only one gene was differentially expressed (**Figure 6A**). This was *NUPR2*, which is related to p53 pathway and DNA damage responses. Thus, we next examined if there were specific changes in the nuclei of the FTD neurons. We found that the shape of the nuclei in the FTD neurons was significantly rounder, indicated by decreased eccentricity, as well as significantly smaller size compared to healthy control neuron nuclei (**Figure 6B**). Interestingly, both C9-HRE and sporadic FTD neurons displayed a significantly higher number of micronuclei, which are small nuclei next to the actual cellular nuclei that are considered to indicate chromosomal instability and caused by incorrectly repaired or unrepaired DNA breaks (**Figure 6C**). In addition, the distribution of larger and smaller sized nuclei was different in FTD neurons than in the control neurons so that a larger portion of the FTD neurons had smaller nuclei (**Figure 6H-J**). When treated with Topotecan, which interacts with topoisomerase I and causes the formation of DNA double-strand breaks, there was a significant increase in the number of γH2A.X-positive foci inside the nuclei in all neurons, indicating that Topotecan increased DNA damage. However, there were no evident differences between the DNA damage response between the FTD neurons as compared to control neurons, nor were there differences in the response between the sporadic and C9-HRE neurons. At baseline, C9-HRE-carrying neurons appeared to display a slightly higher number of γH2A.X foci compared to control neurons, although this difference was not significant (**Figure 6H-J**).

**Figure 6:**
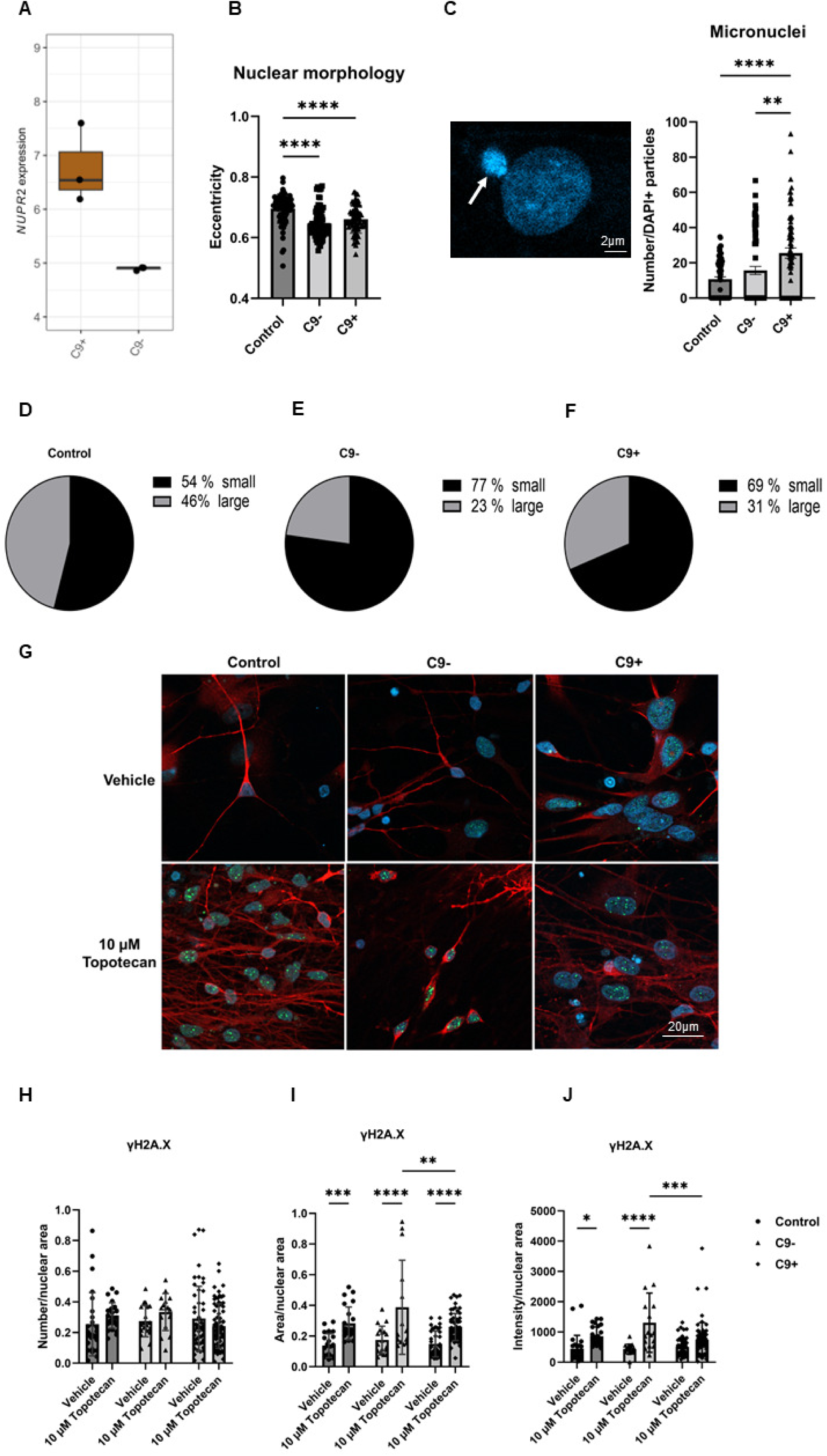
*NUPR2* is the only differentially expressed gene between sporadic and C9-HRE FTD neurons and the neurons display altered nuclear morphologies and presence of micronuclei. *NUPR2,* potentially related to the DNA damage pathway, is the only differentially expressed gene when comparing the C9-HRE and sporadic FTD neurons (**A**). Both sporadic and C9-HRE-carrying FTD neurons display nuclei, which are significantly rounder in shape as well as significantly smaller compared to healthy neuron nuclei, indicated by altered nuclear eccentricity (**B**), and a higher number of micronuclei (**C**). A representative image of a micronucleus next to the nucleus is shown (arrow). Kruskal-Wallis test followed by Dunn’s multiple comparisons test was used. *n* [Control] = 74; *n* [C9-] = 17; *n* [C9+] = 21. In addition, the distribution of larger and smaller sized nuclei is different in FTD neurons than that in control neurons (**D-F**). Topotecan treatment significantly increases the number of γH2A.X-positive foci in the nuclei in all neurons, indicating increased DNA damage. At baseline, C9+ neurons display a slightly higher number of γH2A.X foci compared to HC neurons, although this difference is not statistically significant (**H-J**). Kruskal-Wallis test followed by Dunn’s multiple comparisons test was used. *n* [Control] = 26-28; *n* [C9-] = 15-18; *n* [C9+] = 44-64. Statistically significant differences are shown as *p < 0.05, **p <0.01, and ***p <0.001.

## Discussion

In the present study, the major aim was to contribute to the better understanding of the underlying mechanisms of neurodegeneration in FTD with a special focus on synaptic dysfunction. We investigated whether the synapses are impacted in the FTD patient-derived iPSC neurons and whether the synaptic function and dendritic spine morphologies in C9-HRE and sporadic FTD neurons are similarly or differentially affected.

The C9-HRE leads to several distinct pathogenic mechanisms, including C9orf72 protein loss-of function related to *C9orf72* haploinsufficiency, accumulation of toxic RNA foci, containing the C9-HRE RNA that disturbs RNA metabolism, and accumulation of the harmful DPR proteins [8, 9, 53]. It is suggested that these events occur simultaneously and lead to a cascade of other downstream alterations, including changes in autophagy, stress responses, vesicular trafficking, mitochondrial function, RNA metabolism, synaptic function, and DNA damage [54]. We did not observe differential expression of C9orf72 proteins in C9-HRE-carrying neurons, although they showed a mild but non-significant trend towards lower C9orf72 protein levels compared to control and sporadic FTD neurons. Interestingly, the highest C9orf72 protein levels were detected in the sporadic FTD neurons, which showed significantly higher C9orf72 levels as compared to both control and C9-HRE FTD neurons. The reason for this remains unclear and the underlying mechanisms warrant further examination in future studies. These findings suggest that the C9-HRE-carrying iPSC neurons, at least in the present study, do not show indications of *C9orf72* haploinsufficiency in a similar manner to *e.g.,* patient brain or blood cells. Despite this, the iPSC-derived neurons from C9-HRE carriers displayed the gain-of-toxic-function C9-HRE-associated pathological hallmarks, the RNA foci as well as poly-GP and poly-GA DPR proteins, which were not present in the sporadic FTD or control neurons. Even though previous studies have reported cell toxicity caused by the RNA foci and DPR proteins [55–58], our studies did not validate this. The absence of cell toxicity was confirmed by measurements of secreted NfL levels, a clinically used biomarker of neurodegeneration in patient blood and CSF samples, which indicated undetectable levels of this biomarker in both sporadic and C9-HRE FTD patient neurons. This could be partially due to the missing *C9orf72* haploinsufficiency as it is suggested that haploinsufficiency may potentiate the toxic functions of RNA and DPR accumulations [59]. However, so far, studies have not shown a correlation between DPR pathology and the degree of neurodegeneration or clinical symptoms [60]. It is also suggested that the iPSCs might lose ageing-related characteristics, such as epigenetic modifications including methylation, during cellular reprogramming and, therefore, might not exhibit completely similar pathological features to those observed in patient brains [61]. Additionally, the relatively short culture period of the iPSC neurons might not allow the long-term pathological effects of the C9-HRE to take place.

One of the main pathological alterations in the brain of FTD patients is accumulation of TDP-43 aggregates in cytosolic protein inclusions [5, 6]. Approximately half of all FTD patients display pathological TDP-43 depositions in the brain [62]. Under physiological conditions, TDP-43 shuttles between the nucleus and cytosol, but it is mainly present in the nucleus where it is crucial for RNA regulation, including transcription, alternative splicing, and mRNA stabilization. TDP-43 also partakes in stress granule formation and their maintenance [63, 64]. Under pathological conditions, the shuttling of TDP-43 is disturbed by several post-translational modifications leading to its accumulation and aggregation in the cytoplasm [64]. TDP-43 pathology can be correlated to cognitive decline and neurodegeneration [65] and it occurs downstream of DPR pathology. A previous study has shown that even though the DPR inclusions in the patient brain are typically TDP-43-negative, poly-GP co-localizes with TDP-43 in neurites [66]. We observed that the iPSC-derived neurons from FTD patients, regardless of their C9-HRE status, displayed altered TDP-43 nucleus-to-cytosolic shuttling. The FTD neurons showed a significant increase in the levels of cytoplasmic TDP-43, but no TDP-43-positive cytoplasmic inclusions were detected. This could indicate that alterations in TDP-43 subcellular localization may represent an early feature in FTD pathogenesis, but the accumulation of insoluble pathological TDP-43 inclusions requires time and appears later during the disease. This hypothesis is further supported by other studies showing that disturbed TDP-43 nuclear clearance is sufficient to cause loss of normal TDP-43 function in the neurons without cytoplasmic inclusions present [67, 68]. TDP-43 dysfunction impairs several downstream mechanisms, including protein degradation and autophagy [69]. Several studies show p62/SQSTM1 accumulation in different brain regions in FTD and ALS patients [70] as well as cellular models of TDP-43 pathology [69]. The accumulation of p62 protein is an indicator of defective autophagy [71]. Dysfunctional autophagy and impaired protein clearance may increase the accumulation of TDP-43 aggregates and exacerbate neuronal pathologies. Also, iPSC-derived cortical neurons have previously been reported to show increased p62 levels [72, 73]. Agreeing with these findings, our study also suggests that p62/SQSTM1 accumulates in both the sporadic and the C9-HRE-carrying FTD neurons as indicated by a significant increase in the size and intensity of p62/SQSTM1-positive vesicles. The fact that the number of p62/SQSTM1-positive vesicles was not increased at the same time with the enlarged size and intensity, but rather showed a trend towards a decrease in the FTD neurons, could suggest that the clearance of the p62/SQSTM1-containing vesicles is defective and the vesicles may fuse together. Interestingly, our previous studies in the skin fibroblasts from the same patients, from which the iPSCs were generated for this study, indicated similar results [32]. Thus, autophagy function and protein degradation might be altered in the iPSC-derived neurons from FTD patients, regardless of the C9-HRE status. Whether this is the case can be explored in the future examinations of these neurons.

Besides characterization of the different neuropathologies and their pathological mechanisms, research has focused on structural and functional alterations at synapses and changes in different neurotransmitter systems related to C9-HRE and FTD. Our previous study indicated alterations in dendric branching and spine morphologies as well as compromised neuronal function and altered NMDA receptor function associated with overexpression of the C9-HRE in mouse neurons [31]. Also, studies by us and others have indicated that the C9orf72 proteins as well as poly-GP DPRs locate at synapses [31, 74, 75]. Thus, we assessed dendritic spine number and morphology the iPSC-derived neurons from FTD patients in the present study. The total number of dendritic spines was significantly decreased with a significant reduction especially in the mushroom spines in both C9-HRE and sporadic FTD neurons, suggesting loss of mature, functional spines. In the C9-HRE neurons, these changes were accompanied with a significant increase in the stubby spines, a spine type often associated with neurodegeneration. No changes in the spine head sizes were detected, proposing that the remaining spines are functional and that their strength of synaptic transmission is not altered. These findings suggest that pathological changes in dendritic spine number and morphology and the associated synaptic dysfunction represent common mechanisms underlying both sporadic and C9-HRE-associated FTD.

Functionally, the iPSC neurons derived from sporadic and C9-HRE FTD patients showed a significantly weaker response to GABA stimulus when compared to healthy controls. In addition, a non-significant trend towards diminished responses to glutamate and KCl were detected. Notably, functional changes in the GABAergic and glutamatergic systems have been observed in FTD patient brains [81, 82]. It has been reported that the changes in synaptic transmission correspond with decreased expression of several AMPA and NMDA receptor subunits as well as hyperexcitability in layer V excitatory neurons [83]. Also, it is suggested that neurochemical and functional GABAergic deficits are commonly seen in FTD, especially in bvFTD and PSP [84]. Altogether, our results support the previous findings of impairments in both excitatory and inhibitory systems in FTD patients, which may lead to an imbalanced neurotransmission in their brains. Further supporting synaptic dysfunction, the RNA-seq analysis revealed upregulated DEGs impacting synaptic functions, including synaptic signaling, glutamatergic transmission, and pre- and postsynaptic membrane in the iPSC-derived neurons from FTD patients in comparison to control neurons. Dysregulated synaptic gene expression appeared to be involved in FTD neurons regardless of the C9-HRE carriership. The fact that we observed upregulation instead of downregulation of these synapse-associated genes might implicate potential compensatory mechanisms related to the dysfunctional synaptic neurotransmission in these neurons. Several other studies have reported that the C9-HRE leads to a dysregulation of genes involved in vesicular transport, endosomal trafficking, lysosomal function, proteostasis, DNA damage, synaptic transmission, and neural differentiation [29, 85–88]. One study focused on those changes in C9-HRE carriers without TDP-43 pathology and showed that the C9-HRE is responsible for the alterations in the expression and splicing of genes involved in synaptic vesicle fusion or formation, vesicle transport, endosomal trafficking, chaperone-associated protein aggregation, and DNA repair [85], results that are highly in accordance with our findings in the present study.

Interestingly, the RNA-seq studies revealed only one DEG between the sporadic and C9-HRE FTD neurons. *NUPR2* gene, encoding nuclear protein 2, was upregulated in the C9-HRE-carrying as compared to sporadic FTD neurons. *NUPR2* is a paralogue of *NUPR1*, which is activated in a p53/TP53-dependent manner in response to DNA damage [89]. *NUPR1* is also associated with transcriptomic regulation of mitochondria [90]. It is suggested that ER stress is another factor upregulating the expression of *NUPR1* [91]. In line with these findings, potentially suggesting alterations in the nuclei and DNA damage response, was the significantly altered nuclear morphology observed in the FTD neurons. The nuclei were significantly smaller and rounder in appearance. In addition, we observed a significantly larger number of micronuclei, which are associated with genomic instability leading to drastic DNA re-arrangements [87–89]. However, these changes were detected in both sporadic and C9-HRE FTD neurons, suggesting that the upregulated *NUPR2* expression in the C9-HRE neurons may reflect some other, yet unrecognized, cellular alterations specifically taking place in these neurons. Although we did not observe obvious signs of neurodegeneration in the FTD patient neurons, our findings indicate that they do show pathological changes in the nuclei, which might render the neurons more vulnerable to neurodegeneration if challenged by additional cellular stress. The nuclear alterations might also be related to disturbed DNA damage response. The C9-HRE neurons showed a trend towards a slightly higher number of γH2A.X-positive foci in their nuclei in the untreated baseline conditions, although the difference was not statistically significant. Both the C9-HRE-carrying and sporadic FTD neurons showed significantly increased intensity and size of γH2A.X-positive foci in their nuclei in response to Topotecan, a drug inducing DNA damage. However, the response was similar to the increase in control neurons, suggesting that the FTD neurons do not show clear impairments in their DNA damage response. γH2A.X is phosphorylated upon double-strand break formation, leading to several downstream effects, such as the activation of p53 pathway, which mediates several other cellular responses, such as apoptosis and cell death, cell cycle arrest, or DNA repair [90]. Most interestingly, TDP-43 pathology can be linked to DNA damage and TDP-43 inclusions are suggested to induce p53-dependent apoptosis of iPSC-derived cortical neurons leading to increased neuronal death [91]. A previous study utilizing iPSC-derived brain organoid slice cultures from C9-HRE carriers indicated increased genomic instability and DNA damage in the neurons [87], suggesting that the potential nuclear changes and altered DNA damage response should be further investigated in the iPSC-derived models from FTD patients.

Taken together, our present study shows that the sporadic and C9-HRE-carrying FTD patient-derived iPSC cortical neurons recapitulate the main cellular pathologies and similar structural and functional synaptic disturbances as detected in the patient brains, suggesting imbalanced excitatory and inhibitory neurotransmission. The underlying molecular-level mechanisms may be linked to altered expression and function of several genes regulating synaptic structures and function, which provide specific targets to be investigated in future studies. All in all, our data suggest that the synaptic deficits are a common feature in FTD neurons, regardless of whether they carry the C9-HRE or not and that synaptic dysfunction is an important target for upcoming therapeutic interventions and biomarker development in FTD. Moreover, the present study proposes that FTD patient-derived iPSC neurons represent a valid model for discovering and testing different neuromodulatory approaches aiming at alleviating synaptic dysfunction in FTD.

## Availability of data and materials

The datasets used and/or analyzed during the current study are available from the corresponding author upon reasonable request.

## Acknowledgements

The authors would like to thank University of Eastern Finland Cell and Tissue Imaging Unit (Kuopio, Finland) for providing the infrastructure and help for microscopy and the Biocenter Kuopio National Virus Vector Laboratory at University of Eastern Finland (Kuopio, Finland) for packaging the GFP-AAV vector. Part of the computational analyzes were performed on servers provided by UEF Bioinformatics Center, University of Eastern Finland, Finland, supported by the Biocenter Finland.

## Declarations

### Ethics approval and consent to participate

All the participants gave written informed consent. The research in human subjects is performed in accordance with the ethical standards of Declaration of Helsinki and approved by the Research Ethics Committee of the Northern Savo Hospital District, Kuopio, Finland (ethical permit 16/2013). Studies on FTD patient–derived iPSC-neurons are performed with the permission 123/2016 from the Research Ethics Committee of the Northern Savo Hospital District.

## Authorś contributions

**NH** conceptualization, methodology, investigation, validation of methods, data acquisition, analysis of results, writing original draft, writing review, editing and funding acquisition. **TH:** methodology, validation of methods, analysis. **SH**: investigation, data analysis, review and editing. **AS:** investigation, data acquisition analysis, review and editing. **DH:** investigation, data acquisition, writing-review and editing. **HR:** investigation, methodology, validation of methods, writing-review and editing. **AD:** investigation, data acquisition, analysis, review and editing. **SRN:** investigation, data acquisition, analysis, review and editing. **SK:** data acquisition and analysis, review and editing. **MK:** methodology, validation of methods, review and editing. **EK:** methodology, validation of methods, review and editing. **KP:** methodology, validation of methods, data acquisition, review and editing. **JK:** methodology, validation of methods, data acquisition, review and editing. **TM:** resources, supervision, writing-review and editing. **MT:** methodology, supervision, writing-review and editing. **MH:** conceptualization, funding acquisition, resources, supervision, writing-review and editing. **JKo:** resources, supervision, writing-review and editing. **AMP:** funding acquisition, resources. **ES:** resources, writing-review and editing. **AH:** conceptualization, data curation, writing original draft, writing review, editing, funding acquisition, supervision, project administration, resources

**All authors read and approved the final manuscript.**

